# Oviposition behaviour is not affected by ultraviolet light in a butterfly with sexually-dimorphic expression of a UV-sensitive opsin

**DOI:** 10.1101/2023.01.11.523610

**Authors:** Jose Borrero Malo, Daniel Shane Wright, Caroline Nicole Bacquet, Richard M. Merrill

## Abstract

Animal vision is important for mediating multiple complex behaviours. In *Heliconius* butterflies, vision guides fundamental behaviours such as oviposition, foraging and mate choice. Colour vision in *Heliconius* involves ultraviolet (UV), blue and long-wavelength sensitive photoreceptors (opsins). Additionally, *Heliconius* possess a duplicated UV opsin, and its expression varies widely within the genus. In *Heliconius erato*, opsin expression is sexually dimorphic; only females express both UV-sensitive opsins, enabling UV wavelength discrimination. However, the ecological pressures that have driven these sex-specific differences in visual perception remain unresolved. *Heliconius* females invest heavily in finding hostplants to lay their eggs, a behaviour heavily reliant on visual cues. We tested whether UV vision is used for oviposition in *H. erato* and *Heliconius himera* females by manipulating the availability of UV in behavioural experiments under naturalistic conditions. We found that UV did not influence the number of oviposition attempts or the number of eggs laid. In addition, their hostplant, *Passiflora punctata*, does not reflect UV wavelengths, and models of *H. erato* female vision suggest only minimal stimulation of the UV opsins. Overall, these findings suggest that UV wavelengths do not directly affect the ability of *Heliconius* females to find suitable oviposition sites. Alternatively, UV discrimination could be used in the context of foraging or mate choice, but this remains to be tested.

## Introduction

Animal colour vision mediates a multitude of complex behaviours. Colour vision is achieved via wavelength discrimination (independent of intensity), where the inputs of two or more different photoreceptors (i.e., opsins, which differ in spectral sensitivity) are compared (Kelber, 1999; Kelber & Pfaff, 1999). In most insects, colour vision is based on three photoreceptor classes, encoded by different opsin genes, sensitive to ultraviolet (UV), blue (B) and long-wavelengths (LW) (Briscoe & Chittka, 2001; Kelber, 2006). In contrast, the visual systems of butterflies are highly diverse with differing numbers of photoreceptor classes and sensitivities among families, genera, species and even between sexes within a species (Briscoe, 2008; McCulloch *et al*., 2017; Van Der Kooi *et al*., 2021). Visual system diversification in butterflies and moths has occurred through independent opsin duplication and has been attributed to changes in light availability and habitat use (Sondhi et al. 2021).

Neotropical butterflies of the genus *Heliconius* are a well-studied example of adaptation and speciation (Merrill *et al*., 2015). These butterflies possess striking wing patterns that often converge between two or more distantly related species due to Müllerian mimicry (Müller, 1879). Wing colours are also involved in interspecific and intraspecific communication, and contribute to assortative mating (Jiggins *et al*., 2001; Estrada & Jiggins, 2008; Melo *et al*., 2009; Merrill *et al*., 2014). However, only a few studies have investigated the role of *Heliconius* visual systems from an ecological context (Finkbeiner *et al*., 2017; Dell’Aglio *et al*., 2018), despite vision being largely responsible for mediating these complex behaviours.

Vision plays a crucial role in *Heliconius* behaviour and guides fundamental processes such as intraspecific communication (Estrada & Jiggins, 2008; Merrill *et al*., 2014), foraging (Toure *et al*., 2020) and hostplant selection (Gilbert, 1982). Relative to other butterflies, *Heliconius* have some of the largest brains and invest heavily in the visual neuropile, suggesting selection for well-developed vision (Montgomery *et al*., 2016). In addition to the ultraviolet (*UVRh1*), blue (*BRh*) and long-wavelength (*LWRh*) sensitive opsins found in most insects, *Heliconius* butterflies possess a duplicated ultraviolet opsin (*UVRh2*) (Briscoe *et al*., 2010). Additionally, some *LWRh*-expressing cells possess lateral filtering pigments that shift the spectral sensitivity towards red enabling *Heliconius* to discriminate wavelengths in the long-wavelength range (Zaccardi *et al*., 2006; McCulloch *et al*., 2022).

Within the *Heliconius* genus, opsin expression is variable. Some species, such as *Heliconius melpomene*, have independently lost expression of one of the two UV opsins, with documented pseudogenization events (McCulloch *et al*., 2017). *Heliconius erato*, on the other hand, is sexually dimorphic-males only express *UVRh2* whereas females express *UVRh1* and *UVRh2* (Figure S1)(McCulloch *et al*., 2016, 2017). It is possible that in *H. erato, UVRh1* is located in the W-chromosome such as in *H. charithonia* (Chakraborty *et al*., 2022). A recent study investigated whether *H. erato* was capable of discriminating UV wavelengths: Finkbeiner and Briscoe (2021) tested *H. erato* females (which expresses *UVRh1* and *UVRh2*), *H. erato* males (expresses only *UVRh2*) and *H. melpomene* (both sexes express only *UVRh1*) in a laboratory experiment and found that only *H. erato* females are capable of discriminating between UV wavelengths. However, the ecological pressures that have driven these species-and sex-specific differences in visual perception remain unresolved.

While changes in opsin expression patterns within the *Heliconius* clade are well-documented (McCulloch *et al*., 2016, 2017, 2022; Catalán *et al*., 2019), the adaptive function of these changes in gene expression remains unclear. Understanding the ecological pressures for UV discrimination in *H. erato* females may give us insight as to why expression of this gene varies between and within *Heliconius* species. In contrast to males, female *Heliconius* spend most of their time searching for host plants for oviposition (Benson, 1978), which involves careful visual inspection (Benson, 1978; Brown Jr, 1981). Females also avoid laying on hostplants where other eggs or larvae are present. Indeed, some *Passiflora* species have evolved extra-floral nectaries that reassemble yellow eggs to discourage ovipositing females, but when these egg-mimics are experimentally painted green, females tend to lay more eggs (Williams & Gilbert, 1981). *H. erato* females use leaf shape as an oviposition cue and can learn new shapes, which may explain the plasticity of leaf shape in *Passiflora* (Dell’Aglio *et al*., 2016). The importance of choosing suitable *Passiflora* vines suggests that this behaviour is under strong natural selection (Jiggins, 2017), and evidence suggests that *Heliconius* female vision plays a key role for finding suitable oviposition sites. Therefore, UV wavelength discrimination in *H. erato* females may be an adaptation to facilitate hostplant recognition, but the role of UV vision in this regard has not been tested.

Here, we test whether UV vision is used for oviposition in *H. erato* females and its closely related species *Heliconius himera* by manipulating the availability of UV wavelengths in behavioural experiments. In particular, we address the following questions: (1) Does UV light affect oviposition behaviour in *H. erato cyrbia and H. himera* females? Given that UV is found under natural sunlight conditions, we expect that reducing UV availability will lead to a reduction in the number of eggs laid. (2) Does this behaviour differ between species ? Compared to *H. himera, H. erato* has a larger investment in the visual system (Montgomery & Merrill, 2017), so we predict this effect to be stronger in this species. (3) Are there UV cues in the hostplant and how are these perceived by females ? Given that young shoots are the most nutritious parts of the hostplants and where *H. erato* females preferentially lay eggs (Benson *et al*., 1975; Jiggins, 2017), we predict that UV reflectance will be highest for this part of the hostplant. Finally, we use visual modelling to quantify the stimulation of the *H. erato* visual system by its hostplant *P. punctata*

## Materials and Methods

### Butterfly rearing and maintenance

Wild *Heliconius erat*o cyrbia were caught in forests near Balsas (3°43′60′′S,79°50′45′′W) and *H. himera* near Vilcabamba (4°15′38′′S,79°13′21′′W), in Southern Ecuador. Wild individuals were used to establish stocks at the Universidad Regional Amazónica Ikiam in Tena, Ecuador. Butterfly stocks were kept in outdoor insectaries, in 2 × 2 × 2.3 m cages, fed a 20% sugar solution and had access to pollen from *Lantana sp*. and *Psiguria sp*. flowers. Eggs were collected regularly from hostplants (*Passiflora punctata-*a hostplant of both species (Jiggins *et al*., 1997)), and the larvae were individually reared in pots on fresh leaves from *P. punctata*. For behavioural trials we tested a total of 26 *H. erato cyrbia* and *10 H. himera* females, due to low butterfly stocks.

## Experimental Design

Experiments were conducted between November 2021 and January 2022, under natural sunlight. To manipulate UV in the light environment, two experimental cages (100 × 200 × 235 cm; Supplementary Figure 2) were fitted with either clear UV-blocking (transmission 400-750 nm; LEE #226) or UV-transmitting filter sheets (transmission 300-750 nm; LEE #130). The filters were attached to the top, left and outwards-facing sides of the cage in order to filter the morning sunlight coming from the southeast (Supplementary Figure 2).

The experimental assay lasted six days, during which, a group of females (1-6 individuals) was introduced into each experimental cage. Some individuals (10 *H. erato cyrbia* and 1 *H. himera*) were tested twice (thus, these individuals were tested in 12 trials instead of 6). All females were introduced into the experimental cages at least 24 hours prior to the experiment to acclimate and were confirmed to have laid eggs on a *P. punctata hostplant* that was left in the experimental cage overnight. On the first day of the behavioural experiment, the two cages were randomly assigned a light treatment (UV+ or UV-), thereby controlling for changes in natural sunlight during the assay by testing both treatments in parallel. The following days, each group was tested with the opposite treatment, alternating until both cages had each light treatment (UV+ / UV-) three times under (Figure 1C).

**Figure 1:**
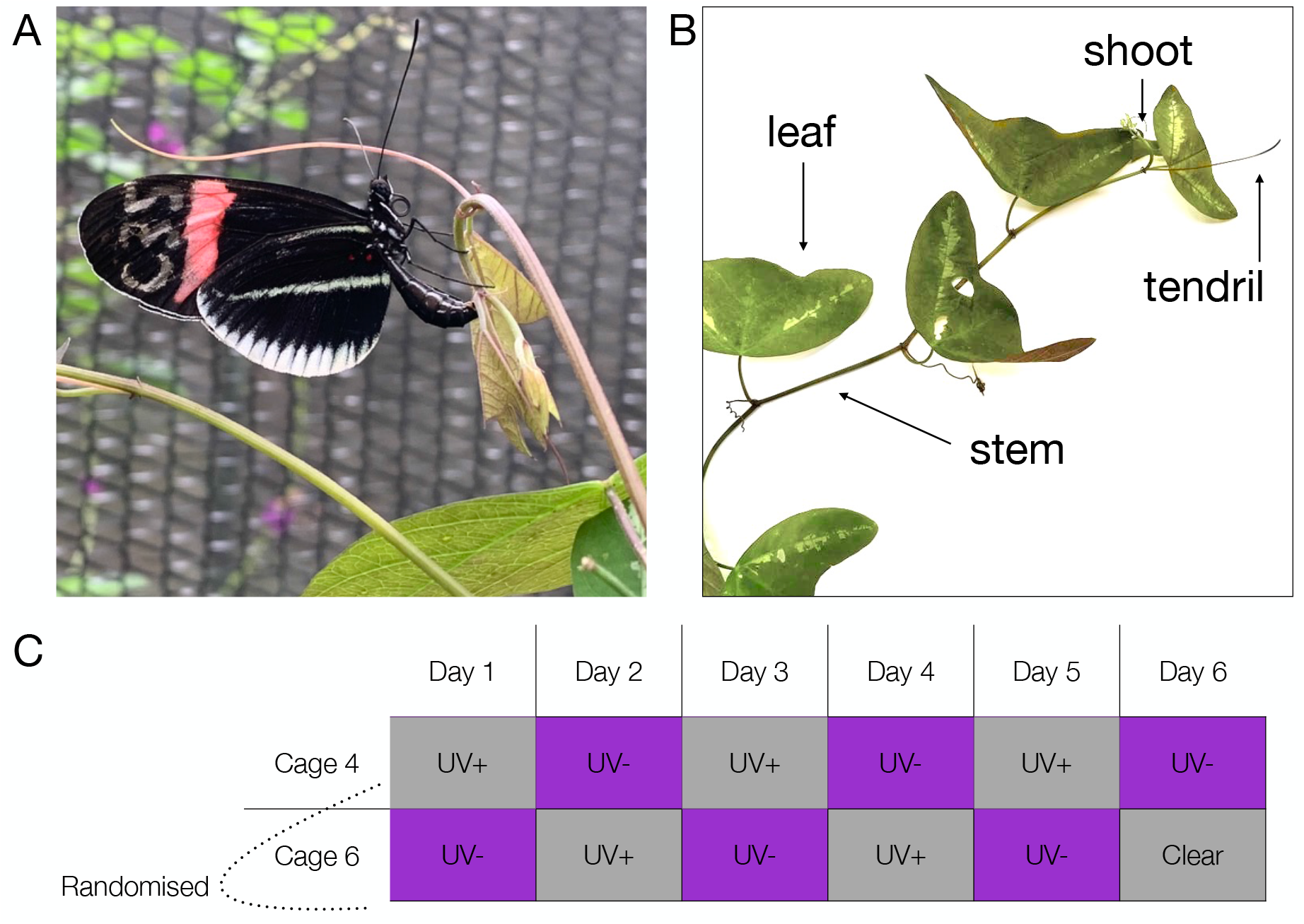
Overview of the experimental setup. (**A**) *Heliconius erato cyrbia* female attempting to lay an egg on a *Passiflora puctata* shoot. (**B**) Parts of hostplant *P. punctata*, where *H. erato* lay eggs. (**C**) Protocol of behavioural experiment.

Before each trial, the filters were fitted to the experimental cages and a *P. puctata* hostplant was placed at the centre of the cage. For each trial, the butterflies were observed for 2h in the morning (between 8:30-12:00). Females were free to fly around the cage, feed on the artificial feeders and a pollen plant, and lay eggs on the plants. For each female, we recorded the number of oviposition attempts, number of eggs laid, and the hostplant “part” (shoot, leaf, tendril, or stem) where the egg was laid (Figure 1B). Oviposition attempts were scored as each time a butterfly landed on a hostplant and moved its abdomen with the ovipositor towards the plant, and each movement of the abdomen towards the plant was counted as an individual oviposition attempt. Egg oviposition after an attempt was scored as an attempt and an egg. The sum of the three trials with the same treatment was combined for analysis.

## Light measurements

Light measurements were taken using a Flame Miniature Spectrometer (Ocean Optics Inc., Dunedin, FL, USA) connected to a UV-VIS optical fibre (P400-2-UV-Vis) with a cosine corrector (Ocean Optics CC-3-UV). In the morning (8:00-12:00), downwelling and side-welling irradiance (in μmol/(m2*s)) was measured in the two experimental cages under the different light treatments (UV+ / UV-). For all measurements, the weather conditions were categorized as sunny (< 50% cloud coverage (cc.)), cloudy (>50% cc.) and overcast (100% cc.). All irradiance measurements were processed and visualized using the pavo 2.2.0 package (Maia *et al*., 2019) in R (R Core Team, 2021).

### Reflectance spectrometry and visual modelling

Reflectance measurements of the hostplant *P. punctata* used were taken using a Flame Miniature Spectrometer connected to a PX-2 xenon light source (spectral range 220nm-750nm) and a UV/Vis reflection probe (Ocean Optics Inc., Dunedin, FL, USA). All reflectance measurements were standardized with a white reflectance standard (Ocean Optics WS-1). For reflectance measurements, the illuminating and reflection probe was placed at a 45° angle at a distance of 1 mm from the plant tissue, and we recorded three measurements per plant (integration time: 2500 milliseconds per scan). All reflectance measurements were then processed and visualized using the pavo 2.2.0 package (Maia *et al*., 2019) in R (R Core Team, 2021). For each plant tissue, 3 biological replicates were measured across 5 individual plants (3 measurements × 3 biological replicates (shoot, stem, leaf) × 5 individuals) = 45 measurements per plant part).

The visual perception of the hostplant was modelled with previously published *H. erato* visual system data (McCulloch *et al*., 2016, 2022) using the pavo 2.2.0 package (Maia *et al*., 2019). For the visual model, we used the following photoreceptor sensitivities of *H. erato* females: *UVRh1* λ_max_355nm, *UVRh2* λ_max_ 390nm, *BRh* λ_max_470nm, *LWRh-green* λ_max_ 555nm and a fifth photoreceptor class *LWRh-red* λ_max_ 590nm that occurs through expression of a red filtering pigment in combination with the green rhodopsin (McCulloch *et al*., 2016, 2022). We then calculated the photoreceptor quantum catch, which estimates the light captured by the visual system (Kelber *et al*., 2003) under each experimental light environment condition (UV+ / UV-) against a green foliage background (Maia *et al*., 2013). The quantum catches were calculated as:

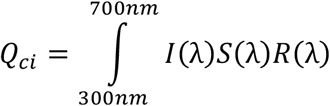

where: I(λ) is the irradiance measured in the experimental light conditions, S(λ) is the reflectance spectrum of the stimulus and R(λ) is the photoreceptor sensitivity based on the equations of (Govardovskii *et al*., 2000; Hart & Vorobyev, 2005).

## Statistical analysis

All statistical analyses were conducted in R (R Core Team, 2021) and plots were created with the *ggplot2* package (Wickham, 2011) (see supplementary material for script in R Markdown). We fitted generalized linear-mixed models (GLMM) with the *glmer* function in the lme4 package (Bates *et al*., 2015) to test if oviposition behaviour was affected by the presence or absence of UV, and tested how the number of oviposition attempts and/or eggs laid was influenced by the fixed effects (and their interactions): (i) treatment (UV+ / UV-), (ii) weather (< 50% cloud coverage / >50% cc. / 100% cc.) and (iii) species (erato / himera). Where GLMMs with Poisson distribution were overdispersed, we fitted negative binomial models with the *glmer*.*nb* function in the lme4 package. To avoid pseudoreplication (individuals were tested multiple times), individual id was included as random factors. The random effect structure of the full models was selected based on Akaike comparisons, choosing the model with the lowest AIC value (ΔAIC > 4; (Sakamoto *et al*., 1986; Burnham & Anderson, 2004). Stepwise model reduction of the fixed effects based on statistical significance (Crawley, 2002) was then conducted using likelihood ratio tests (LRT) via the *drop1* function to identify the minimum adequate statistical models. To estimate the parameters of significant fixed effects, we used parametric bootstrapping (nsim= 1000, pbkrtest package (Halekoh & Højsgaard, 2014)). For fixed effects with more than two categories (e.g., weather), we conducted pairwise comparisons using post hoc Tukey corrections with the emmeans package (Lenth *et al*., 2019).

## Results

### UV does not affect oviposition behaviour

The availability of UV wavelengths did not significantly affect the number of oviposition attempts (LRT=0.0855, df=1, p= 0.39) (Figure 2A). Similarly, there were no species differences in the number of oviposition attempts (LRT=0.1459, df=1, p= 0.72). Neither the UV treatment (LRT=1.6258, df=1, p= 0.20) (Figure 2B) nor species identity (LRT=1.0624, df=1, p= 0.31) had a significant effect on the number of eggs laid. The number of oviposition attempts significantly differed by weather (LRT= 21.764, df=2, p= 0.001). Post-hoc pairwise comparisons indicated that females had fewer attempts on days with full cloud coverage than on sunny days (Z=-2.837 p=0.0127). Weather also had a significant effect on the number of laid eggs (LRT=11.641, df=2, p= 0.004); more eggs were laid on sunny days (<50% cc.) than on overcast (100% cc.) days (Z=-2.446, p=0.038) (Supplementary Figure 4).

**Figure 2:**
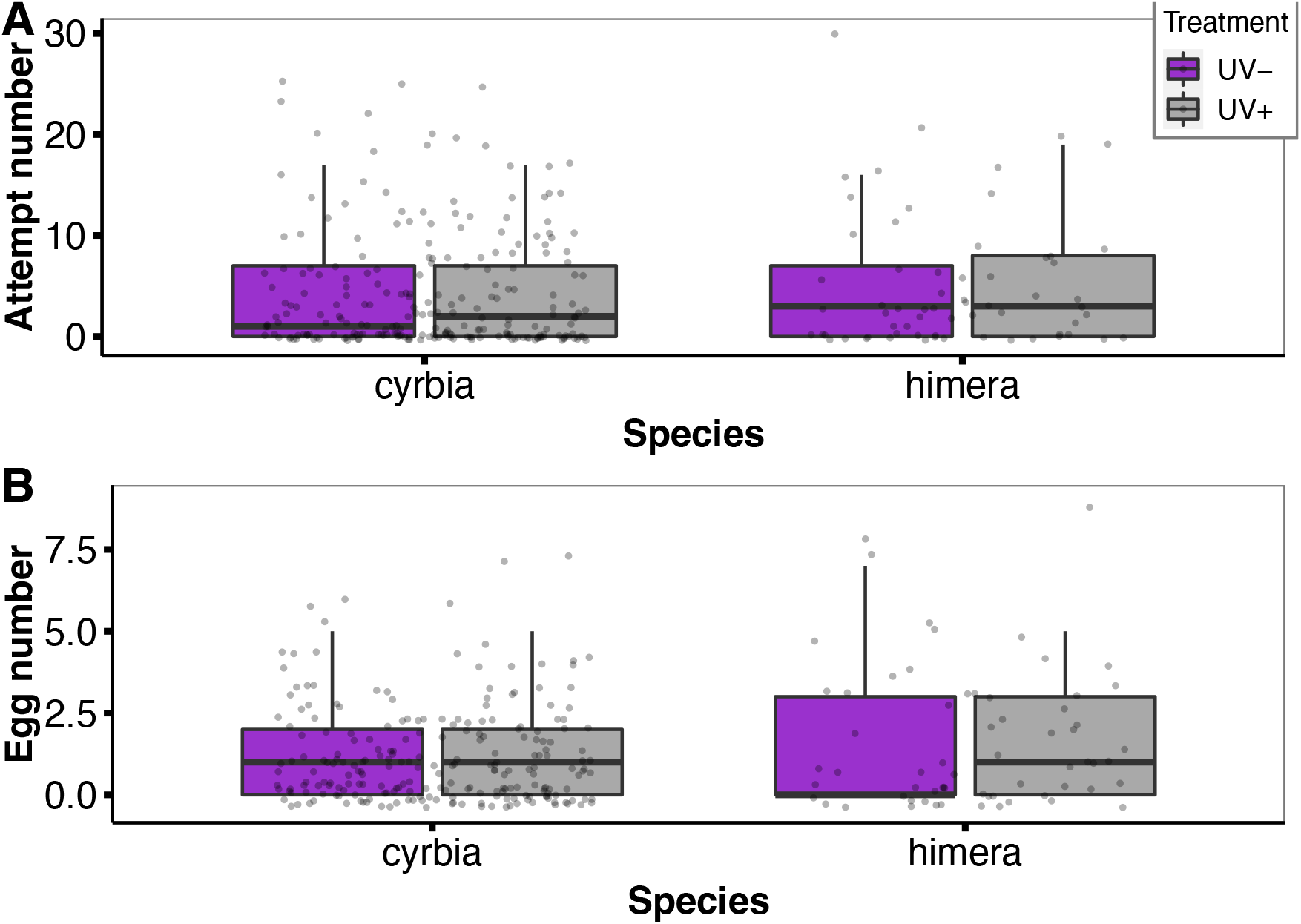
UV manipulation did not affect the (**A**) number of oviposition attempts or (**B**) the number of eggs laid on the hostplant. Grey boxes represent the number of attempts/eggs in the control treatment (UV+) and purple boxes represent the number of attempts/eggs in the UV-light environment. Error bars represent ± 1 standard error.

### Females prefer to lay eggs on shoots

The number of oviposition attempts significantly differed by plant part (LRT=57.164, df=3, p< 0.001) (**Error! Reference source not found**.A). Post-hoc tests showed more attempts on shoots compared to leaves (Z=5.268, p <.0001), stems (Z=5.988 p <.0001) and tendrils (Z= 5.506, p <.0001). The number of eggs significantly differed by plant part (LRT=24.704, df=3, p< 0.001) (Figure 3B), but this was not influenced by the UV treatments (the *treatment:plant-part* interaction was non-significant; X^2^= 0.7731, p=0.85588). As with the number of eggs, post-hoc analyses revealed that more eggs were laid on the shoots compared to leaves (Z=4.85, p <0.0001), stems (Z=2.780, p= 0.03) and tendrils (Z=2.654, p = 0. 04).

**Figure 3:**
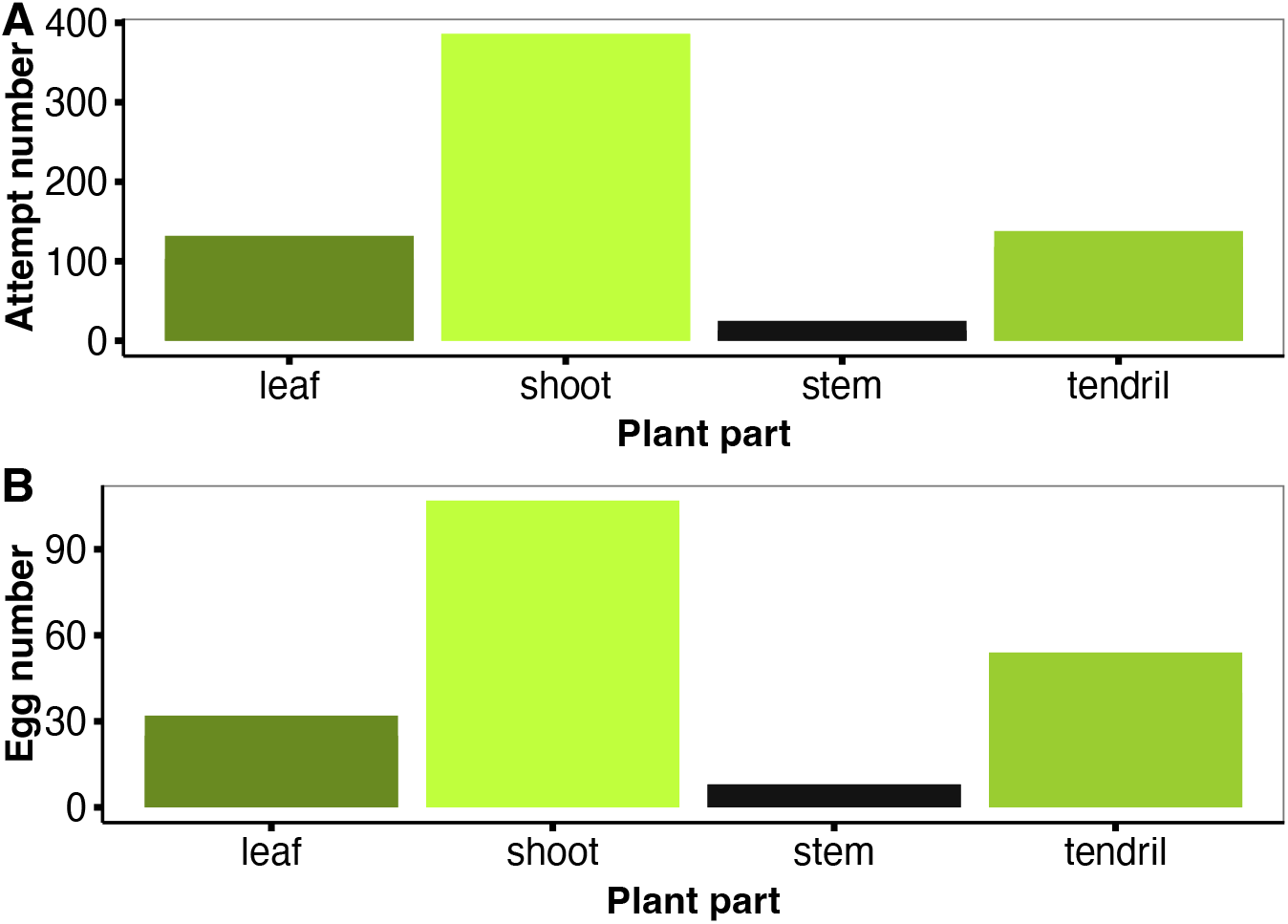
Total number of (**A**) oviposition attempts and (**B**) eggs per plant part, throughout the behavioural experiment. (**C**) Parts of hostplant *P. punctata*, where *Heliconius* lay eggs

### Hostplant does not reflect UV and visual models show little-to-no stimulation of the UV photoreceptors

The spectral reflectance curves of different parts (shoot, stem, leaf, and white patches on the leaf) of the hostplant *P. punctata* are presented in Figure 4A. The observed reflectance curves are likely due to the presence of light-absorbing chlorophyll in all plant parts (Chappelle *et al*., 1992); reflectance peaks are present at ∼ 550 nm and >680nm, and there is low reflectance below 500nm, with very little reflectance in the UV range (300-400 nm).

**Figure 4:**
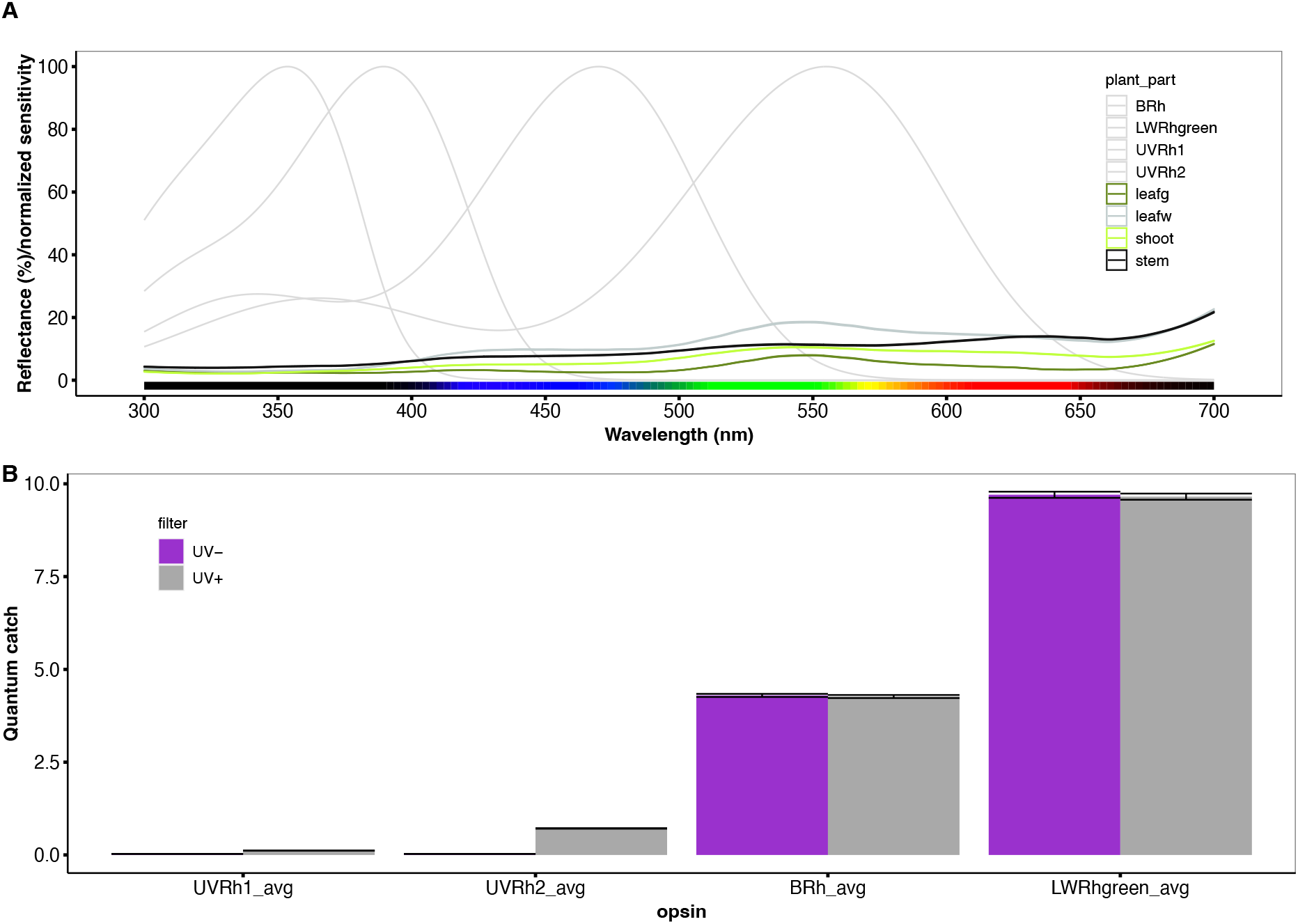
(**A**) Reflectance spectra of *P. punctata* parts. Grey lines on background indicate the normalized spectral sensitivities of *H. erato*. **(B**) Quantum catch estimates of the female *H. erato* visual system when viewing the shoots of *P. punctata* against a green foliage background. Quantum catches were calculated for each opsin *UVRh1, UVRh2, BRh1 and LWRh* including the red ‘receptor’. Error bars represent ±standard error.

To estimate visual perception of the host plant by females in the UV-manipulated treatments, we calculated the photoreceptor quantum catches for the shoots of the hostplant-the part where most eggs were laid-against a green foliage background under each experimental condition (UV+ / UV-) (Figure 4B). Under natural sunlight (UV+), *UVRh2* was minimally stimulated, while our models predicted that *UVRh1* was not stimulated. In UV-absent conditions (UV-), neither UV opsin (*UVRh1* or *UVRh2*) was stimulated. In contrast, the blue opsin (*BRh*) and the long-wavelength opsin with and without filtering pigments *(LWRh-green and LWRh-red)* were similarly stimulated under both lighting conditions. The long-wavelength receptor with red filtering pigments (*LWRh-red*) had the highest quantum catch, followed closely by the long-wavelength opsin *(LWRh-green)* (Figure 4B).

## Discussion

Vision plays a crucial role in *Heliconius* behaviours, including mate choice (Estrada & Jiggins, 2008; Merrill *et al*., 2014), foraging (Toure *et al*., 2020) and hostplant selection (Gilbert, 1982). Through a gene duplication event at the base of the *Heliconius* genus, these butterflies have gained a secondary UV-sensitive opsin (Briscoe *et al*., 2010; Bybee *et al*., 2012). However, expression of this opsin varies both between and within species. For example, numerous species have independently lost expression of one of the two UV opsins, with documented pseudogenization events (McCulloch *et al*., 2017). In *Heliconius erato*, UV opsin expression is sexually dimorphic: females express both UV opsins, whereas males only express one (McCulloch *et al*., 2016).

In a recent laboratory study, male and female *H. erato* differed in UV wavelength discrimination (Finkbeiner & Briscoe, 2021). However, the ecological pressures that have driven these sex-specific differences in visual perception remain unresolved. Given the differences in life history between male and female *Heliconius* butterflies, we predicted that UV wavelength discrimination might benefit female-specific behaviours such as oviposition. A strong body of evidence suggests the importance of visual cues for finding suitable hostplants for oviposition in *Heliconius* females. However, our experiments suggest that UV wavelength discrimination in *H. erato* females is not an adaptation associated with oviposition behaviours.

In our experiments, the availability of UV light did not influence the number of oviposition attempts or the number of eggs laid by *Heliconius* females. Our spectral reflectance measures of the hostplant *P. punctata* provides a possible explanation for these results. Overall, we found only minimal UV reflection in any of the hostplant parts (shoot, leaf, or stem) of *P. puctata* where female butterflies laid eggs (Figure 4A). This result is in line with the fact that UV reflectance is usually – but not always – low on leaves (Archetti *et al*., 2009), whereas, in contrast, many flowers reflect UV (Arnold *et al*., 2008). We also used female *H. erato*-specific opsin sensitivities (McCulloch *et al*., 2016, 2022) to estimate the photoreceptor quantum catches when viewing the shoots of *P. punctata-*where most eggs were laid-in both light conditions (UV+ & UV-). Neither UV photoreceptor (*UVRh1* or *UVRh2*) was stimulated under the UV-absent conditions, and under natural sunlight (UV-present), only *UVRh2* was minimally stimulated (Figure 4B). The absence of UV reflectance in the hostplant and little-to-no stimulation of the UV photoreceptors suggests that UV discrimination does not directly affect *Heliconius* female oviposition. However, it is important to note that these conclusions are based on estimates of visual system stimulation which are inherently limited (Dell’Aglio *et al*., 2018; Drewniak *et al*., 2020; Finkbeiner & Briscoe, 2021), further highlighting the importance of our behavioural studies.

The circuitry required for UV discrimination is metabolically costly and may have trade-offs with other components of colour vision (McCulloch *et al*., 2016). Our experiments make an important contribution by ruling out the possibility that UV perception in *H. erato* females is used during oviposition. One alternative is that *H. erato* females may use UV discrimination to detect previously laid eggs. Because of cannibalism in *Heliconius* larvae, females avoid ovipositing in the presence of conspecific eggs on the hostplants. However, neither *H. erato* eggs nor *Passiflora* egg-mimics reflect wavelengths in the UV range (300-400 nm) (Finkbeiner & Briscoe, 2021) so this explanation seems unlikely.

Another possibility is that UV wavelength discrimination is used in female mate choice. Studies in other butterfly groups, such as *Colias* and *Eurema*, have shown that UV reflectance is used by females for conspecific recognition and mate choice (Silberglied & Taylor, 1973; Kemp, 2008). In *Heliconius*, UV opsin duplication co-occurred with the evolution of a yellow pigment (3-hydroxyDL-kynurenine) that reflects UV (Briscoe *et al*., 2010; Bybee *et al*., 2012). However, there are populations of *H. erato* which do not show these yellow patterns, and it is currently unknown whether variation in UV vision exists between populations. Experiments have shown that both male and female *H. erato* individuals prefer to approach UV+ over UV-models (Finkbeiner *et al*., 2017); however, *Heliconius* females do not generally approach males to solicit mating, and these experiments cannot distinguish between UV-guided mating preference behaviours, or more general attraction to UV reflecting cues, which are common in flowers used by these butterflies (see below). Other experiments have manipulated UV reflectance on the wings of *H. erato* and its co-mimic *H. melpomene*, by applying UV-blocking sunscreen, and have found that *H. erato* males more often approached *H. melpomene* females when the UV signal was blocked (Dell’Aglio *et al*., 2018).

However, this does not explain the sexual dimorphism in UV opsin expression in these species. Nevertheless, visual modelling does suggest that female *H. erato* may be able to distinguish between the yellow colours of *H. erato* and *H. melpomene* (Dell’Aglio *et al*., 2018), so although there is little evidence that wing colours play a role in female mate choice in *Heliconius*, it remains an intriguing hypothesis.

A more likely alternative function of UV discrimination in *H. erato* females could relate to foraging. Many insects that forage on flowers, such as bees and butterflies, can perceive UV (Briscoe & Chittka, 2001). Analysis of the reflectance of *Psychotria* and *Psiguria* – two pollen plants used by *H. erato* – found a UV component on the reflectance spectrum of their flowers (Finkbeiner & Briscoe, 2021). Female *Heliconius* have higher nutrient requirements than males, due to egg production, and may need to invest more in foraging for pollen resources. In particular, in *H. charathonia*, which also has sexually dimorphic vision (McCulloch *et al*., 2017), females have been shown to collect significantly more pollen than males (Mendoza-Cuenca & Macías-Ordóñez, 2005). Using a similar experimental design as the one used in the present study, future research could investigate the function of UV discrimination in the context of foraging.

An important caveat of our study is that we used individuals from populations collected from the wild on the western slopes of the Andes in southern Ecuador. Previous studies, which reveal evidence of sexually dimorphic expression of the UV-opsins, used *H. erato petiverana* individuals supplied from a breeder based in Costa Rica (McCulloch *et al*., 2016, 2017). The same subspecies, *H. e. petiverana*, and supplier was also used for laboratory based UV wavelength discrimination experiments (Finkbeiner & Briscoe, 2021). The most recent common ancestor of the *H. erato* clade dates to 200,000-500,000 years ago and since then, over 15 *H. erato* races with different wing patterns have evolved (Van Belleghem *et al*., 2017). Gene expression evolution can be rapid, and especially in visual systems (Seehausen *et al*., 1997; Nandamuri *et al*., 2017). Therefore, it is possible that different *H. erato* populations might differ in their opsin expression patterns, though this has not yet been explored.

To our knowledge, this is one of the very few studies (but see also Veen *et al*., 2017), that have used filters to modify natural sunlight conditions in a behavioural experiment Using natural sunlight conditions as opposed to standardized artificial lighting has advantages and disadvantages. First, sunlight is likely to better represent the lighting conditions found in the habitats of these butterfly species and may elicit more natural behaviour. However, when conducting experiments under natural sunlight conditions, there is considerable variation in the light intensity (see Supplementary Figure 2). Thus, an unintentional difference in light intensity may affect the results. Indeed, weather significantly affected oviposition attempts and the number of eggs laid in our study (Supplementary Figure 4). Specifically, compared to overcast weather, butterflies made more attempts and laid more eggs on sunny days. For this reason, the majority of behavioural studies that have manipulated the presence of UV in the environment using UV-blocking filters have used standardized artificial lighting (Lewis *et al*., 2000; Greenwood *et al*., 2002; Honkavaara *et al*., 2008; Hiermes *et al*., 2021). Nevertheless, under natural conditions-particularly in rainforests-light intensity varies rapidly (Endler, 1993), which will be better reflected by experiments manipulating wavelength under more natural conditions such as ours.

In conclusion, *Heliconius* colour vision plays a crucial role during fundamental behaviours, including mate choice, oviposition and foraging. In contrast to *H. erato* males, *H. erato* females express two UV-sensitive opsins and can discriminate between UV wavelengths, but the ecological significance of this sexual dimorphism remains unresolved. Further research is required to better understand the evolutionary processes that have sex-specific differences in visual perception in *Heliconius*.

Nevertheless, by manipulating the light environment under naturalistic conditions, we have made an important contribution by ruling out UV perception in *H. erato* females as an adaption relating to oviposition behaviours.

## Acknowledgements

We are grateful to the Universidad Reginal Amazónica Ikiam for support in Ecuador. We thank Lucie Queste and Sophie Smith for thoughtful discussions and for assistance in the insectaries. We also thank Anderson Yumbo and Mariana Silva who contributed to data collection. This research was funded by a ERC Starting Grant (851040) to R.M.M.

## Supplementary material

**Supplementary Figure 1:**
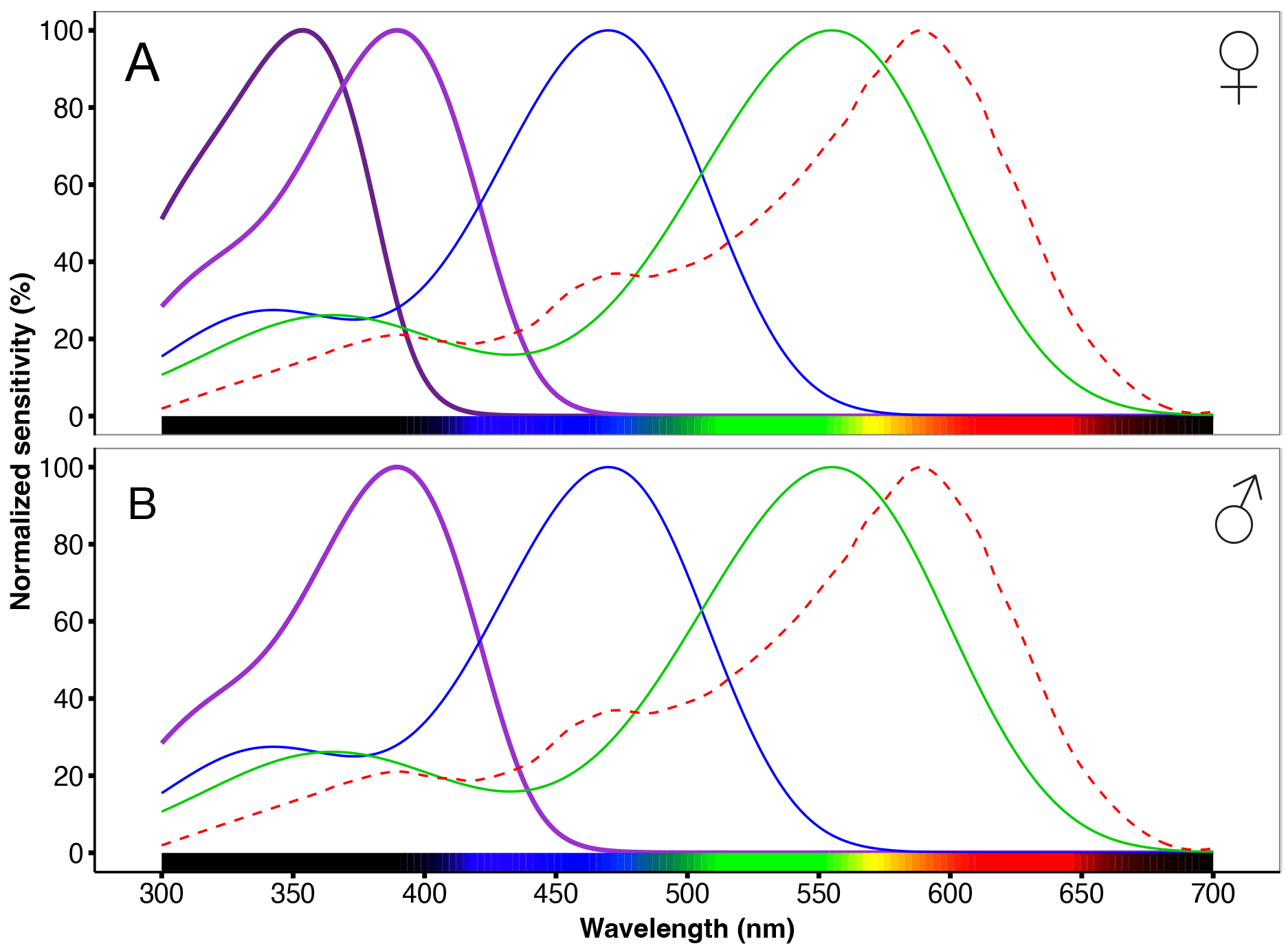
Normalized spectral sensitivities of *Heliconius erato*. (B) Males possess opsins with peak sensitivities: UV 390 nm (*UVRh2*), blue 470 nm (*BRh*) and long-wavelength (green) 555 nm (*LWRh*) (A) Females possess an additional UV opsin (*UVRh1*) that has a peak sensitivity at 355 nm. Additionally, *H. erato* possesses a fifth receptor class, with a peak at ∼ 590 nm due to filtering of the green rhodopsin by a red filtering pigment (red dashed line, from McCulloch et al. 2021).

**Supplementary Figure 2:**
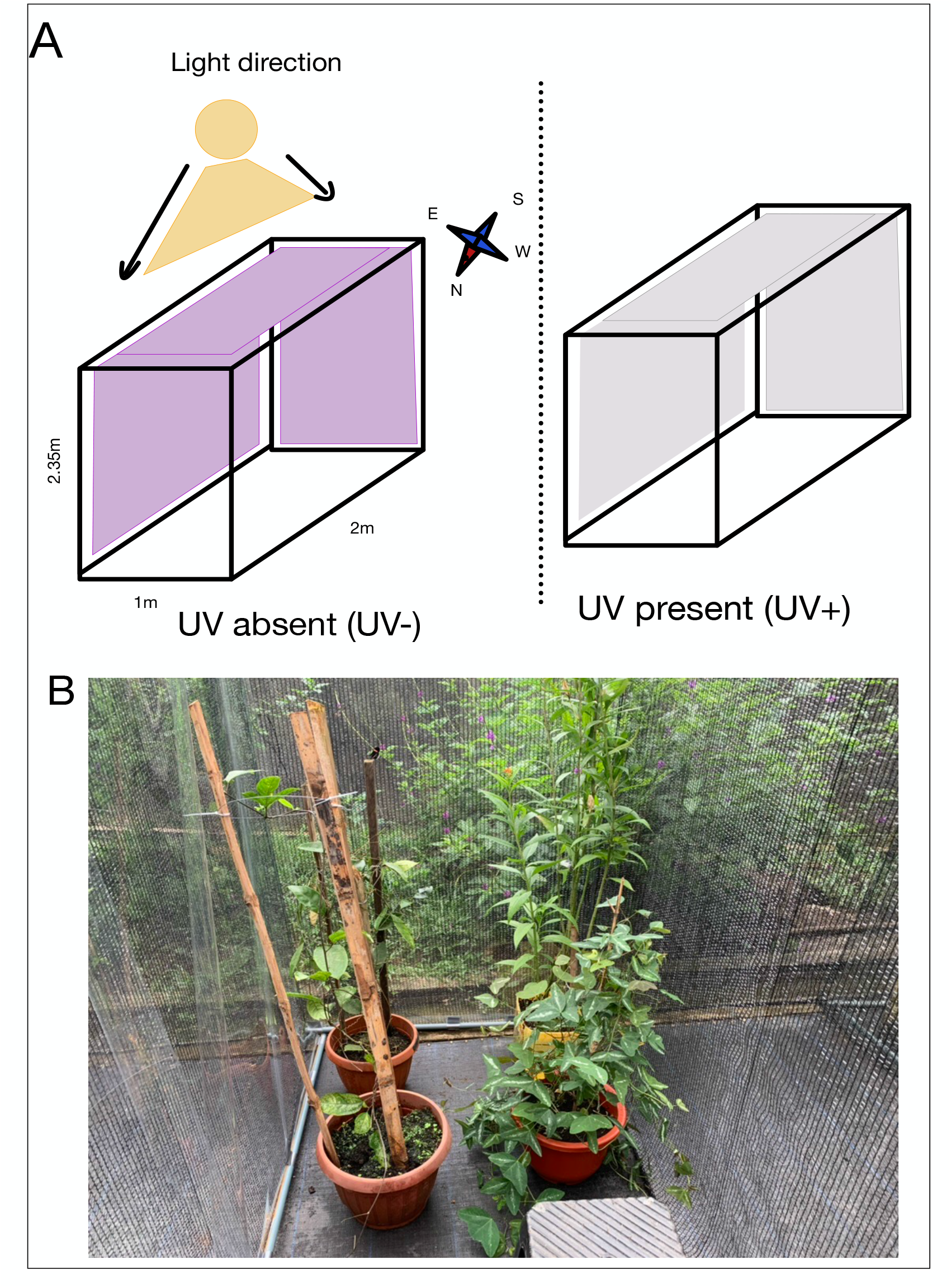
Experimental cage set up. (A) Schematic view of the cage used for the behavioural experiment. Cage on the top left shows experimental cage under UV-absent conditions when fitted with a UV-blocking filter. Cage on the top right shows experimental cage under UV+ present conditions, when fitted with a clear filter. Filters were placed on the east and south-ward facing sides of the cage which received most of the incoming natural sunlight. (B) Picture of the experimental cage fitted with UV-blocking filter with a *Passiflora punctata* hostplant.

**Supplementary Figure 3:**
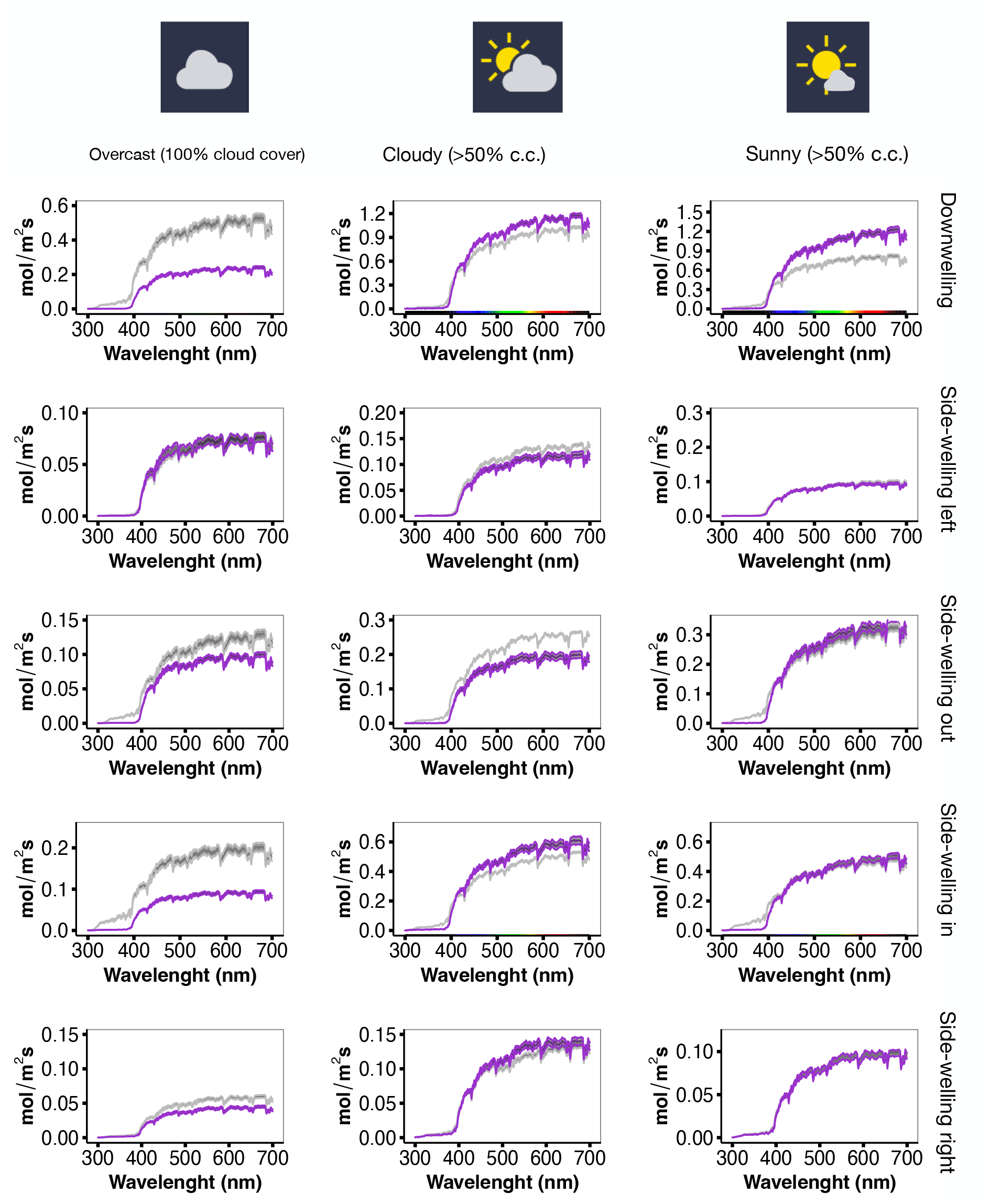
Experimental light irradiance in the UV-Vis range in the under different weather conditions. Purple line represents irradiance in cage with UV-blocking filter UV-absent, grey line represents irradiance in the cage with clear filter UV-present. Shaded areas represent ± one standard error. Left column shows the irradiance spectra under overcast conditions (100% cloud coverage), middle column shows the irradiance spectra under cloudy conditions (>50% c.c.) and right column shows the irradiance under sunny conditions (<50 % c.c.). Top three rows show sides of the cage fitted with filters (1) downwelling, (2) side-welling out (southeast facing) (3) side-welling left (east facing). Bottom two rows show sides of the cages that were not fitted with filters (4) side-welling in (north facing) & sidewelling right (northwest facing).

**Supplementary Figure 4:**
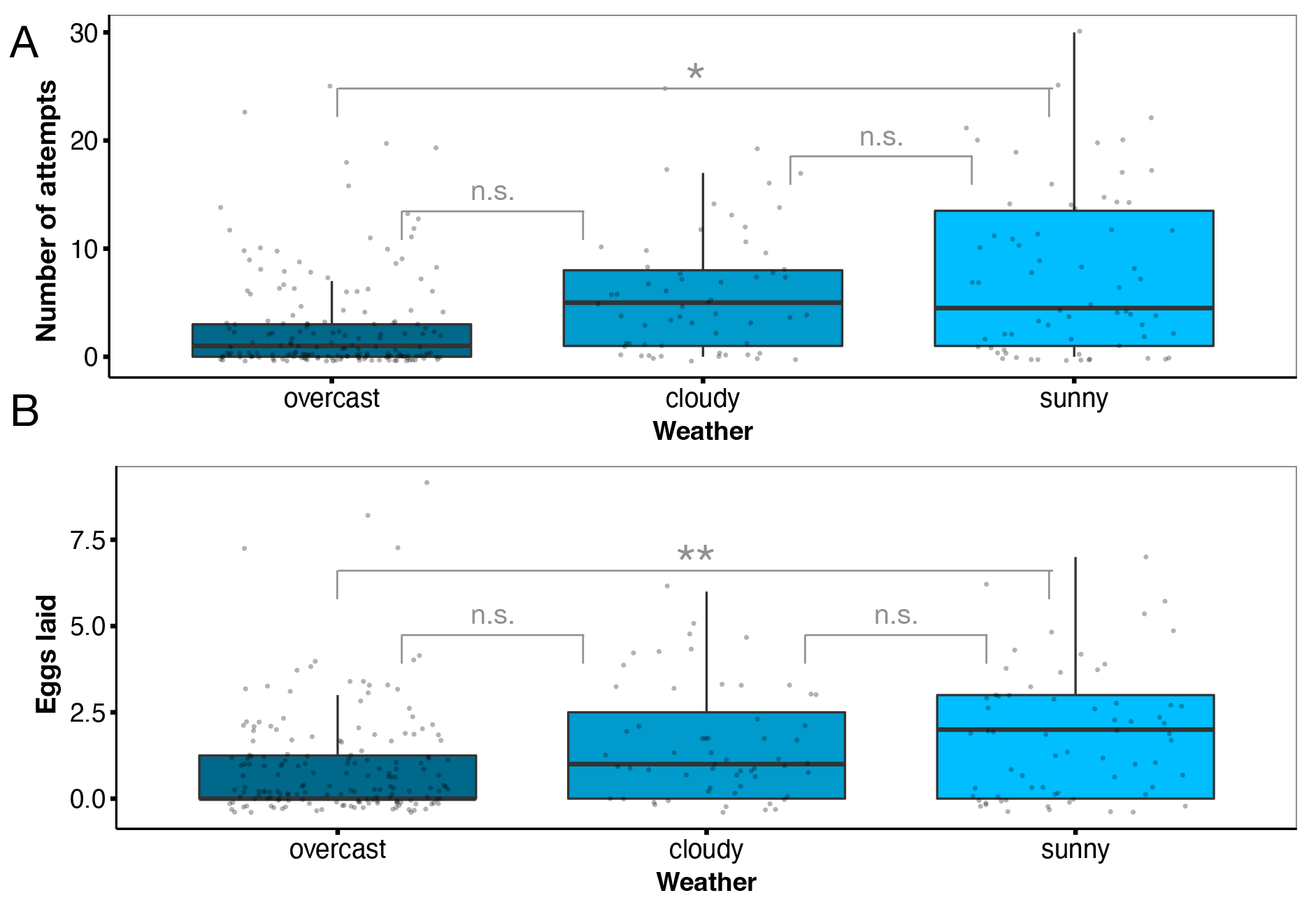
(**A**) Number of oviposition attempts per weather category (**B**) Number of eggs laid per weather category.

**Supplementary Figure 5:**
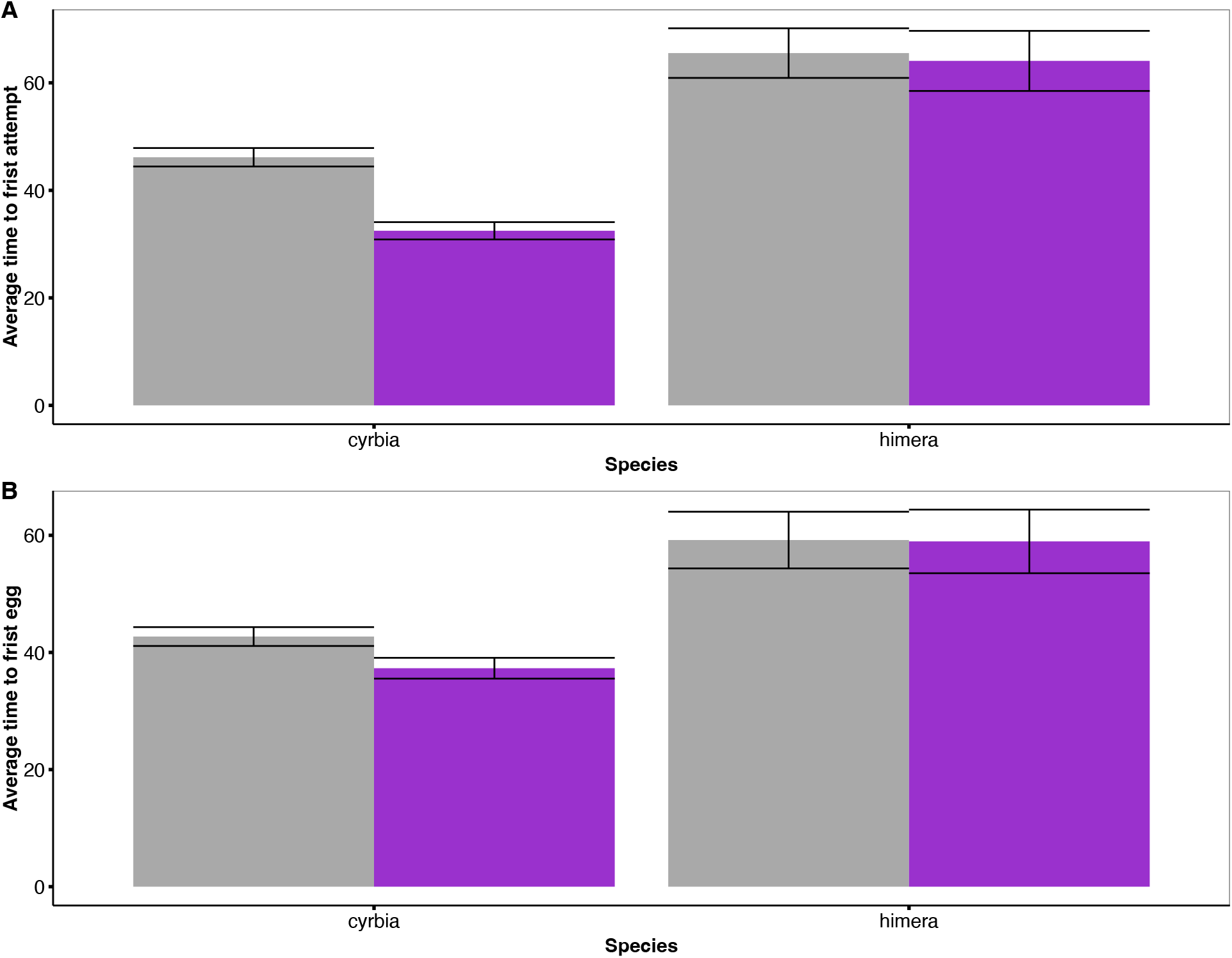
Average time to first (A) oviposition attempt (B) egg laid. Grey bars represent number of attempts/eggs in the control treatment (UV+) and purple boxes represent the number of attempts/eggs in the UV-light environment.

**Supplementary Figure 6:**
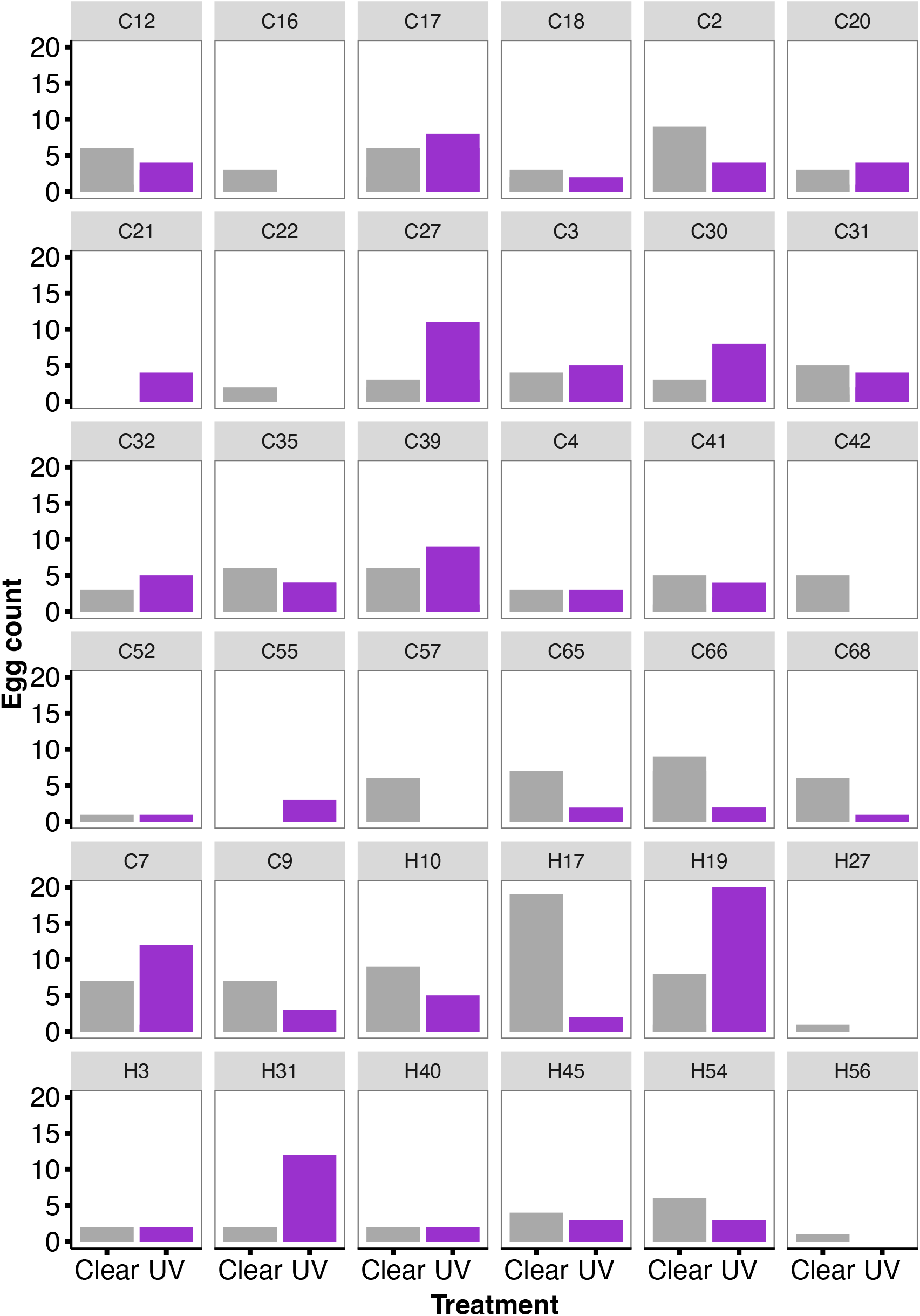
Egg count per treatment by individual. Females marked with C represent *H. erato cyrbia* while *H*. himera individuals are represented with an H.

